# Chromatin-loop extrusion and chromatin unknotting

**DOI:** 10.1101/411629

**Authors:** Dusan Racko, Fabrizio Benedetti, Dimos Goundaroulis, Andrzej Stasiak

**Affiliations:** Center for Integrative Genomics, University of Lausanne, 1015 Lausanne, Switzerland; SIB Swiss Institute of Bioinformatics, 1015 Lausanne, Switzerland; Polymer Institute of the Slovak Academy of Sciences, 842 36 Bratislava, Slovakia

**Keywords:** Chromatin, DNA topoisomerases, DNA knots, Chromatin-loop extrusion, Cohesin

## Abstract

It has been a puzzle how decondensed interphase chromosomes remain essentially unknotted. The natural expectation is that in the presence of type II DNA topoisomerases that permit passages of double-stranded DNA regions through each other, all chromosomes should reach the state of topological equilibrium. The topological equilibrium in highly crowded interphase chromosomes forming chromosome territories would result in formation of highly knotted chromatin fibres. However, Chromosome Conformation Capture (3C) methods revealed that the decay of contacts with the genomic distance in interphase chromosomes is practically the same as in the crumpled globule state that is formed when long polymers condense without formation of any knots. To remove knots from highly crowded chromatin, one would need an active process that should not only provide the energy to move the system from the state of topological equilibrium but also guide topoisomerase-mediated passages in such a way that knots would be efficiently unknotted instead of making the knots even more complex. We show here that the process of chromatin-loop extrusion is ideally suited to actively unknot chromatin fibres in interphase chromosomes.

**SIGNIFICANCE STATEMENT:** Similar to earphone cables crammed into a pocket, long and crowded chromatin fibres that form chromosomes in living cells have the natural tendency to get knotted. This is exacerbated by the action of DNA topoisomerases that transiently cut some chromatin fibres and let other to pass through. Yet, the knotting frequency of chromatin fibres is very low and it has been a puzzle how this is achieved. Recently a novel active mechanism known as chromatin loop extrusion has been proposed to be involved in shaping chromosomes by forming sequential arrays of ca 1 Mb large chromatin loops. Using numerical simulations, we show here that chromatin loop extrusion is ideally suited to remove knots from chromatin fibres.

## INTRODUCTION

Type II DNA topoisomerases that pass DNA segments through each other are required to facilitate various DNA transactions and they are critically important for the segregation of freshly replicated DNA molecules (1, 2). However, DNA-DNA passages occurring in highly crowded DNA molecules in living cells inadvertently lead to formation of knotted DNA (3, 4). Long-lasting knots on DNA are undesirable as tightened DNA knots hinder RNA transcription (5) and lead to DNA breakage (6). A “proof-reading” mechanism is needed to actively remove accidentally introduced knots. There are strong indications that an active knot-removing mechanism operates in highly crowded nuclei of eukaryotic cells. Chromosome conformation capture studies revealed that chromatin fibres in interphase chromosomes have the scaling behaviour characteristic of a fractal globule, which is a condensed polymer state that is free of knots (7). This paucity of knots in chromatin, in addition to avoiding problems created by tightened DNA knots, allows unimpeded decondensation of chromosomal regions where genes are expressed or where replication occurs (8, 9). However, active unknotting mechanisms provide significant conceptual challenges (10). Topoisomerases involved in unknotting would need to be guided to act on those juxtapositions of chromatin fibres where a passage simplifies or removes a knot rather than creates a knot or makes an existing knot more complex (11). Currently, it is unknown how DNA topoisomerases are guided to unknot chromatin fibres. However, numerous recent experimental studies concluded that interphase chromosomes are shaped by an active process of chromatin loop extrusion (12-14). Here we show, using molecular dynamics simulations, that chromatin-loop extrusion process is ideally suited to unknot and decatenate crowded chromatin fibres as well as to segregate chromosomal territories from each other.

### Results and Discussion

It is now broadly accepted that topologically associating domains (TADs) in interphase chromosomes of mammalian cells are formed by chromatin loop extrusion mechanism (15, 16). During chromatin loop extrusion cohesin rings load on chromatin fibres in such a way that they initially form local, small chromatin loops, which are then progressively enlarged by active sliding of cohesin rings along the looped chromatin fibre (15, 16). Until now chromatin loop extrusion was only seen as a mechanism ensuring that border elements of TADs approach each other, resulting in formation of loop-like chromatin regions with increased frequency of internal contacts (12-14, 17, 18) and it was not considered that chromatin loop extrusion can be involved in active unknotting of chromatin fibres.

To investigate whether chromatin loop extrusion can be involved in chromatin unknotting, we performed coarse-grained molecular dynamics simulations of chromatin loop extrusion involving knotted chromatin fibres (see Material and Methods for more detailed description of the simulation procedure). In our proof of principle simulations, we started with knotted chromatin fibres forming closed loops. The closure of loops is justified by the fact that TADs spend significant fraction of time as closed chromatin loops (12). In addition, loop closure prevents the introduced knots from escaping. Figure 1 shows snapshots from a simulation of a loop extrusion occurring in knotted chromatin fibre that approximates a loop-forming TAD (12). The simulated chromatin fibre forms a trefoil knot, which is most frequently observed knot forming in chromatin rings *in vivo* (3). Simulated chromatin loops contain a region mimicking functional elements of the TAD border. There are beads mimicking the presence of bound CTCF proteins, which stop the progression of cohesin rings (12-14). There is also a short region with reduced excluded volume potential that permits passages of other portions of modelled chromatin fibres through that region. Having such a region in modelled chromatin fibres allows us to mimic the action of DNA topoisomerase TopIIB that is known to be bound near CTCF and ensures that there is the possibility of chromatin fibres passing through each other (19). Although type II DNA topoisomerase can permit intersegmental chromatin passages occurring also at other places, the majority of Topo II-mediated passages occur at borders of TADs (20) and that is why we introduced the possibility of Topo II-mediated passages occurring there. After a short thermal equilibration, the process of loop extrusion is started by progressively threading chromatin fibres through the cohesin handcuffs (green-coloured in Fig. 1) and thus progressively enlarging the loop spanned by the cohesin handcuff (Fig. 1b). Simulations show that as the loop extrusion progresses cohesin handcuffs push before them the entanglements forming a knot and therefore the extruded part (brick-coloured), which is behind cohesin handcuffs, is free from knots, whereas the remaining part (blue) contains the original knot. As the loop extrusion proceeds, the knot gets pushed towards the TAD border, where it is unknotted by a passage involving the region with a reduced excluded volume potential (Fig. 1c-d). In our simulations, we initiated chromatin loop extrusion at a position that is equally distant from the left and the right TAD border. In a biological setting this may be a rare situation. However, when loop extrusion starts from any other site and the cohesin ring that first reaches the TAD border is stopped by bound CTCF the second cohesin ring still progresses (14), therefore, the end result is the same, i.e. the knot is pushed towards TAD border and is unknotted there. Simulations aimed to reproduce experimental contact maps (13, 14) and experiments investigating the dynamics of cohesin and CTCF *in vivo* (21) indicate that chromatin loop extrusion occurs repeatedly, i.e. cohesin handcuffs reach the border of TADs and stay there for some time but eventually dissociate and can start a new round. Therefore, all knots that may spontaneously form in the extruded chromatin loop as result of Topo II action assisting transcription or replication will be again unknotted by the next round of chromatin loop extrusion.

**Figure 1.**
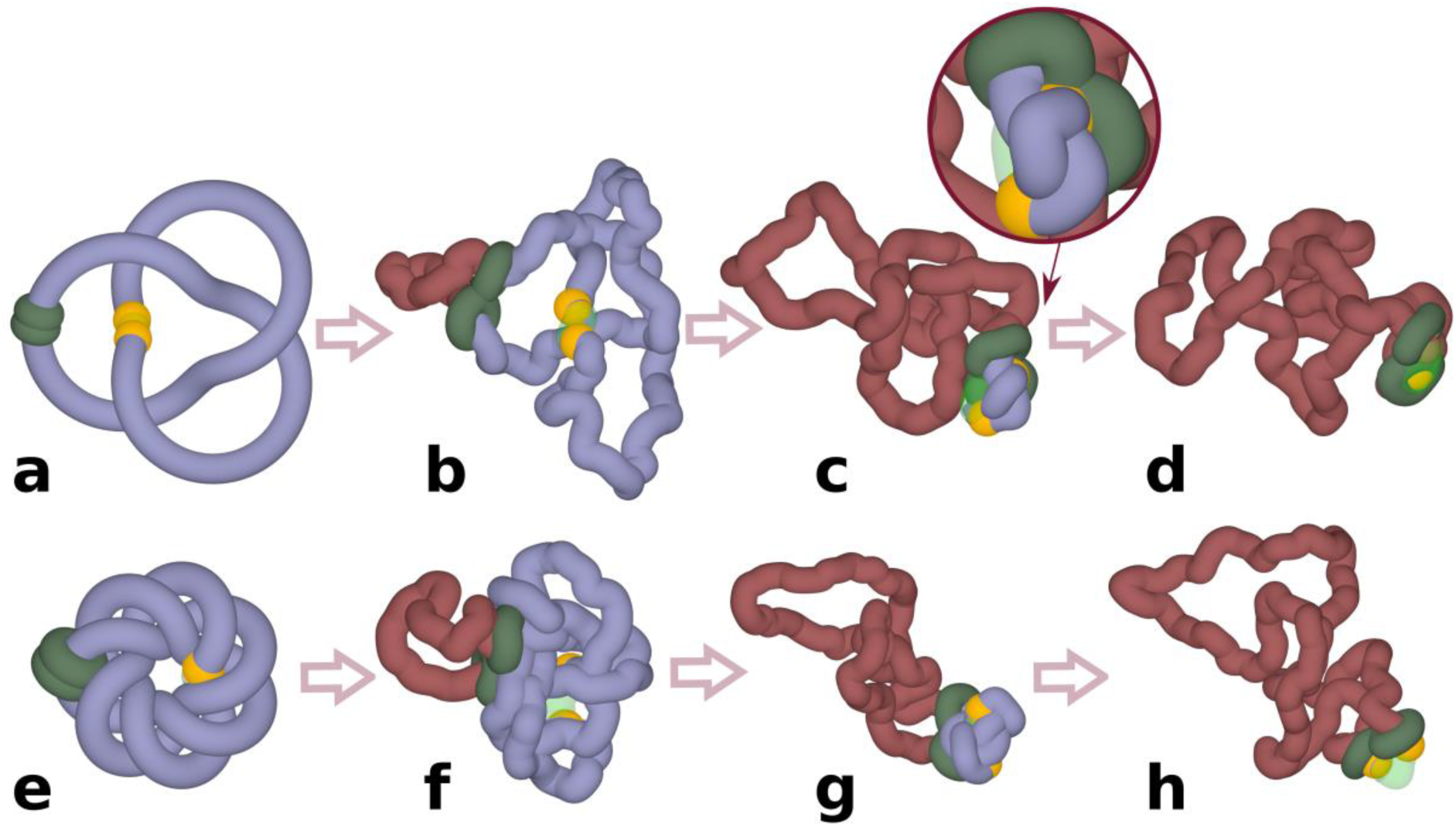
Chromatin loop extrusion unknots individual loop-forming TADs modelled as chromatin rings. a-d. Unknotting of a simple trefoil knot. **e-h**. Unknotting of a complex knot with 14 crossings. **c** and **d** show highly compressed knots being pushed towards region with reduced excluded volume potential, which in our simulations mimic the action of TopIIB known to be present near TADs borders. Inset in **c** shows a close-up of a strongly confined knot, which is about to get unknotted by an intersegmental passage involving a region with reduced excluded volume potential. **d** and **h** show that at the end of simulations the knots are already unknotted. In our simulations cohesin rings (green) are initially arbitrarily placed opposite to borders of modelled TADs. Once simulations are started, stacked rings of cohesin take the form of a quasi-planar handcuff and actively translocate in that form. Borders of TADs are recognizable here by yellow coloured beads that in our simulations stop the progression of cohesin rings and thus mimic the biological action of CTCF proteins located at borders of TADs.

Next, we wanted to test what happens when a more complex knot forms on a chromatin fibre. Since complex torus knots can be formed as products of site-specific recombination (22), we investigated how chromatin loop extrusion could unknot such a complex knots. Figure 1e- h shows loop extrusion occurring in a TAD containing a complex torus knot with 14 crossings. As in the case of the trefoil knot, this more complicated knot gets also unknotted. This indicates that, in principle, chromatin loop extrusion is capable to promote unknotting of even very complex knots.

Simulations presented in Figure 1 were of simplified systems where individual TADs were modelled as independent chromatin loops. In reality, TADs are positioned one after the other along the chromosome and are just separated by short “spacers” (12). In addition, chromatin knots do not need to be very local and confined within individual TADs but can be larger, involving chromatin regions larger than one TAD. To model such a situation, we constructed a system where a chromatin portion containing three TADs was closed into a circle forming a large trefoil knot. Closure into a circle is needed to maintain the knot in this relatively short portion of modelled chromosome. In real chromosomes with thousands of TADs, closure would not be needed to keep the knots from slipping over the ends. Figure 2 shows what happens in such simulations where chromatin loop extrusion occurs in each of the constituent TADs. In order to better visualize the process in these proof of principle simulations, chromatin fibres were not equilibrated before chromatin loop has started. This approach allows us to initially maintain the quasi-symmetric configuration of modelled chromatin portion with three TADs, which is a remnant of the starting configuration. This setting of the simulation makes it easier to see that entanglements are excluded from extruded portions of TADs and accumulate in the inter-TAD portions, which are known to be sites where Top2B is located (19). As already mentioned, the activity of Top2B is accounted for in our simulations by placing short regions with reduced excluded volume potential. As chromatin loop extrusion progresses, the entanglements of knots concentrate in the vicinity of regions with reduced self-avoidance and unknotting occurs at these regions (Fig. 2d).

**Figure 2.**
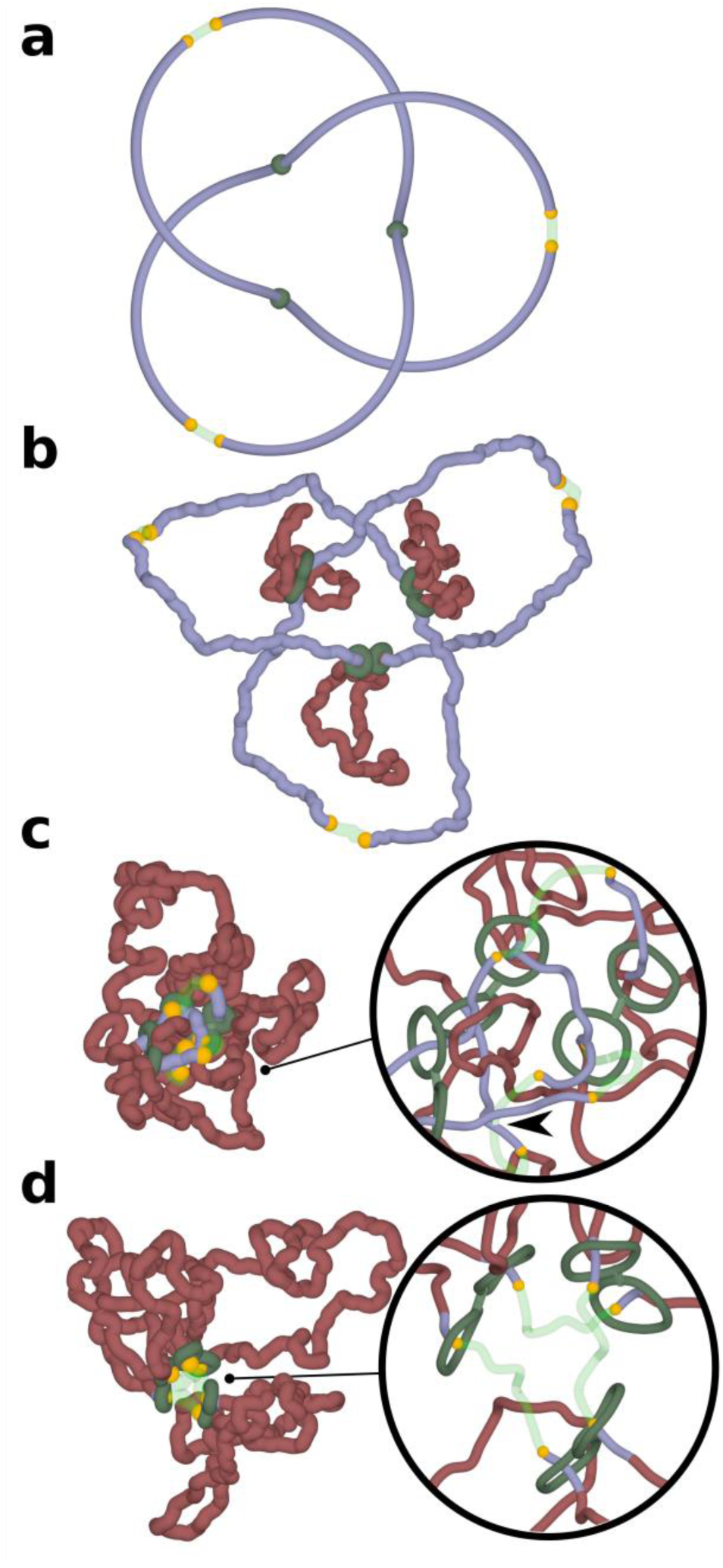
Chromatin loop extrusion unknots a delocalized trefoil knot spanning a modelled chromosome portion containing three TADs. Three TADs, each with two cohesin rings forming handcuffs (green), beads mimicking action of CTCF (yellow) and TopII sites (green semi-transparent) are modelled as presented in Fig.1**. a**. The starting configuration shows that the formed trefoil knot is not localized within an individual TAD but diffuses over three TADs. **b-d**. Chromatin loop extrusion proceeds in all three TADs. The knot becomes excluded from the extruded chromatin portions and is concentrated in the chromatin portions with TopII sites of action, which are modelled as regions with reduced excluded volume potential. As the knot becomes more confined it is more likely to interact with topoisomerases localized just outside of TADs and is eventually unknotted there. Insets show close-ups of regions near TADs borders. For better visibility, the diameter of modelled chromatin fibres and cohesin handcuffs is decreased. In inset shown in **c** the arrow indicates an unknotting passage involving a region with reduced excluded volume potential (semi-transparent) and a chromatin portion that did not pass yet through cohesin rings (blue). In inset shown in **d** the extrusion is nearly finished as cohesin rings are approaching sites with bound CTCF (yellow). The portion of chromatin, which will not be extruded further is already free from knots.

The simulations shown in Figs. 1 and 2 were of relatively simple systems where simulated TADs were not affected by crowding resulting from high concentration of chromatin fibres in chromosome territories (23). To account for the crowding and the large length/diameter ratio of chromatin fibres forming TADs, we performed simulations of highly concentrated long chromatin rings undergoing loop extrusion. Our simulated system was composed of 8 chromatin rings having each size of about 1.6 Mb. Individual rings thus had sizes of large TADs. The concentration of modelled chromatin fibres was set to 30% by adequately adjusting the radius of the confining sphere where these 8 chromatin rings were placed. 30% volume occupation corresponds to the concentration of chromatin in nuclei of eukaryotic cells (24). Subsequently, the system with 8 confined rings was topologically equilibrated by switching off the excluded volume potential and thus permitting thermally fluctuating chains to freely pass through themselves and through each other. After that step, we introduced excluded volume potential acting within and between all 8 chains. At the end of this procedure each modelled chromatin ring was highly knotted (Fig. 3) and each ring was also catenated with at least 3 other rings. The step of producing topological equilibrium is only to test what would happen if one could start chromatin loop extrusion from topologically equilibrated situation. As already mentioned, Hi-C studies established that chromatin in interphase cells is essentially unknotted (7) and studies of chromosome territories in interphase cells showed only very limited intermingling between neighbouring chromosome territories (25) thus indicating the absence of extensive catenation.

**Figure 3.**
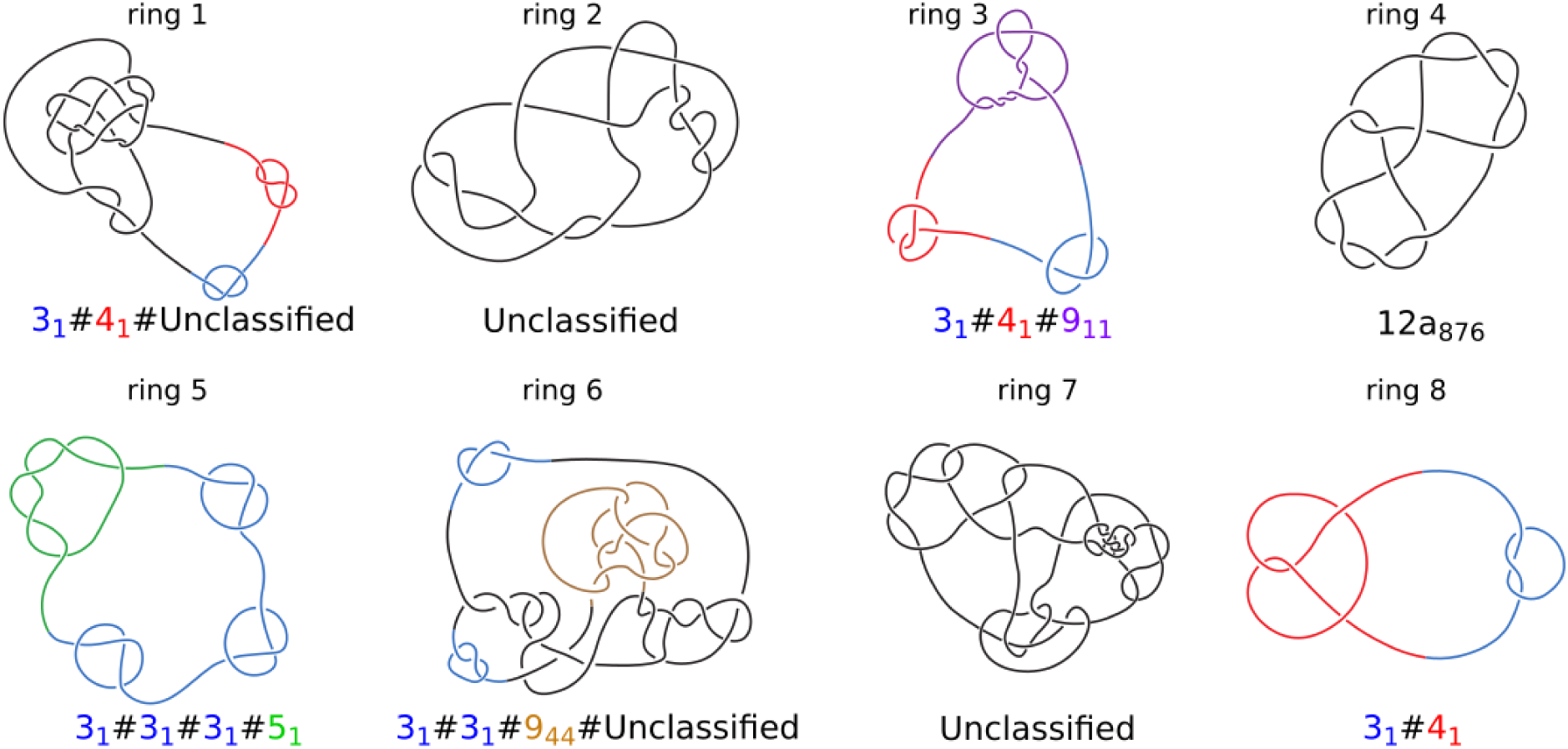
Topology of knotted rings formed upon topological equilibration of 8 modelled chromatin rings. Knot types of all 8 rings forming a system presented in Figure 4. Alexander-Briggs notation of knots uses two numbers to indicate the knot type. The first number written in normal font indicates the minimal number of crossings of a given knot and the second number written as a subscript indicates the tabular position of this knot among the knots with the same minimal crossing number. So for example, 9_11_ indicates a knot that in tables of knots is placed at 11^t^^h^ position among knots which minimal crossing number is 9. The # sign indicates that a knot is formed by a composition or merging together of simpler (prime) knots that are listed as its components. For prime knots with a large number of minimal crossings, e.g. 12, as it is the case of ring 4, the knots are additionally divided into these with alternating pattern of crossings (indicated as a) and with non-alternating pattern of crossings (indicated as n). Standard tables of knots do not include knots with more than 12 crossings and we indicate these knots as unclassified.

Starting from this topologically very complex system composed of 8 knotted and catenated with each other rings that are in addition crowded (see Fig. 4 a), we initiated simultaneous loop extrusion in each of the 8 rings. In each ring the point where the loop extrusion started was the most distant from the region with reduced extruded volume that was used to mimic the presence of Top2B at borders of TADs (19). Panels a and b in Figure 4 show simulated system just before the loop extrusion has started (a) and just after the loop extrusion was completed (b). Each of the simulated rings is presented in a different color. Importantly, although the system started as a mixture of highly knotted and catenated rings, by the end of loop extrusion each ring was both unknotted and decatenated from all other rings. By comparing panels a and b one gets an impression that loop extrusion process demixes individual rings from each other and segregates territories occupied by each of the rings. To quantitatively measure the extent of demixing, we have used a Voronoi space tessellation approach (26) to subdivide the volume of the sphere containing the 8 chains into polyhedra enclosing each bead forming the 8 modelled chromatin rings (see methods section for more detail). We subsequently calculated the area of interface between all Voronoi polyhedra containing beads belonging to different chains. Figure 4 c shows the surface of interface between Voronoi polyhedra centred at beads of one individual chain and these centred at beads of other chains. We call these types of surfaces Voronoi envelopes. Figures 3d and e show the total surface of all Voronoi envelopes for the system before and after chromatin loop extrusion. The graph in Figure 4 f shows how the interfacial area between all Voronoi envelopes decreases during chromatin loop extrusion. Therefore, we can conclude that chromatin loop extrusion not only unknots and decatenates individual rings but also segregates modelled chromatin rings and decreases their intermingling.

**Figure 4.**
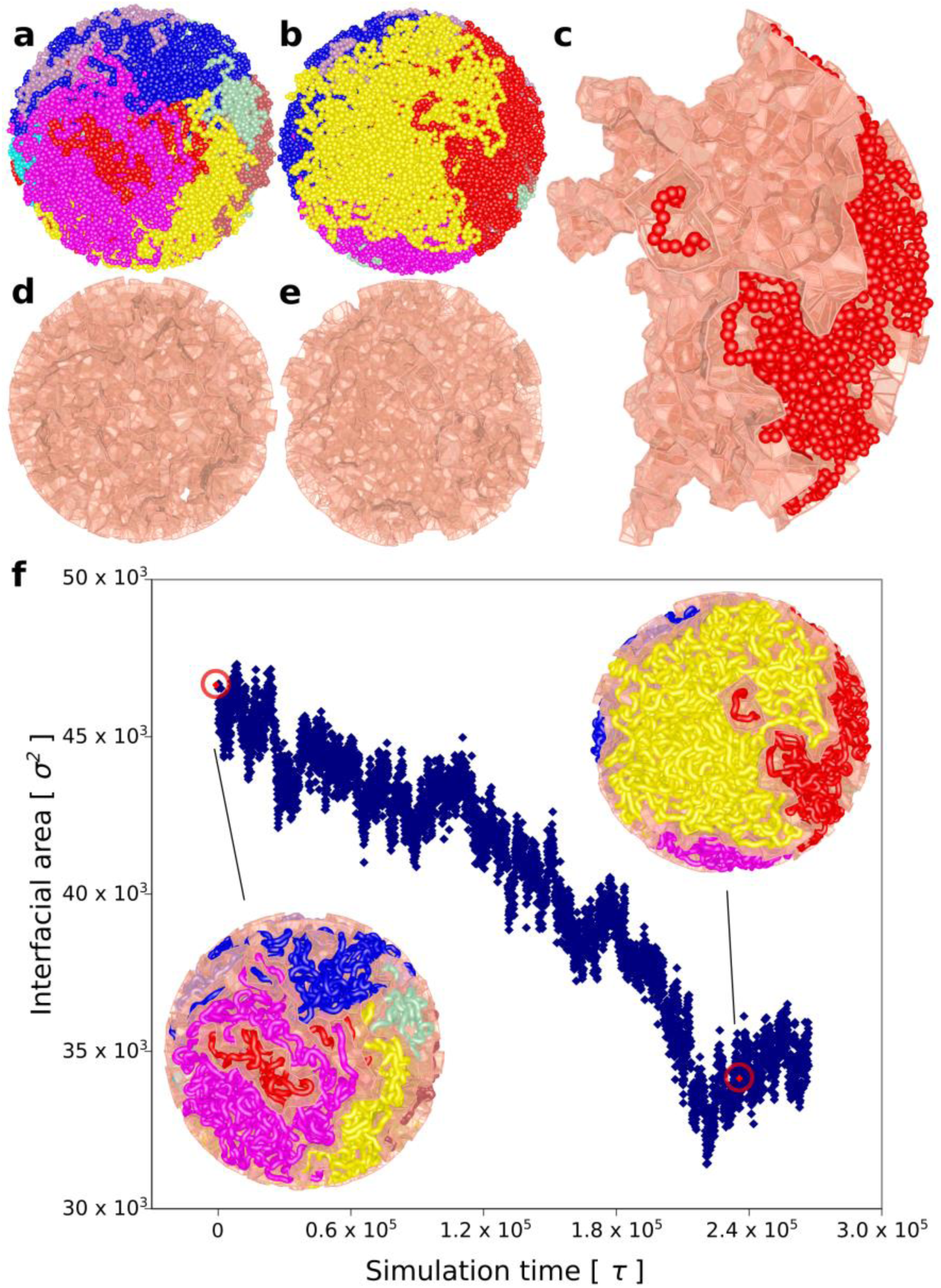
Chromatin loop extrusion demixes topologically equilibrated chromatin loops. a. A snapshot of topologically equilibrated system composed of 8 large chromatin circles (each shown in different colour) that are knotted and catenated with each other. **b**. A snapshot showing how the simulated system changes after each of the rings has undergone chromatin loop extrusion. **c**. One of the 8 chromatin circles is shown together with its Voronoi envelope marking the interface of that chromatin circle with other circles. **d** and **e**. Voronoi envelopes marking interfacial area between different chromatin circles before and after chromatin loop, respectively. **f**. As chromatin loop extrusion progresses the interfacial area between all modelled loops decreases. The snapshots present the confined chains together with their Voronoi envelopes at the moments when chromatin loop extrusion has started and finished, respectively. Red, encircled points indicate the interfacial area measures for the shown snapshots.

Experimental studies of chromosomal territories revealed their limited intermingling (25, 27). It has been a question of how this limited intermingling is achieved in the presence of DNA topoisomerases (28). Uncontrolled action of DNA topoisomerases is expected to result in a highly-intermingled state of chromatin fibres from all decondensed chromosomes during the interphase stage of a cell cycle (7). We showed here that chromatin loop extrusion is likely to be the driving force for unknotting, decatention and segregation of chromatin fibres forming different chromosomes. It is important to add here that unknotting, decatenation and segregation investigated here just depend on the progressive movement of cohesin rings towards TAD borders. That movement might be driven by still undemonstrated translocation activity of cohesin (16) or by transcription-induced supercoiling (29).

It needs to be stressed here that the mechanism of chromatin unknotting, decatenation and demixing by chromatin loop extrusion is very much different from the mechanism of chromatin condensation needed for the formation of mitotic chromosomes. Chromatin condensation mediated by the action of condensin helps to disentangle sister chromatids from each other but this is due to effective shortening of chromatids changing them from long, thin and floppy filaments that easily entangle with each other to short, thick and rigid rods that are unable to entangle with each other (30, 31). In contrast to condensin-mediated condensation of chromosomes, cohesin-mediated chromatin loop extrusion does not induce condensation of loops that passed through cohesin rings but just moves chromatin fibres through cohesin rings.

It is a popular saying that one can’t unscramble an egg. Apparently, chromatin loop extrusion process can very well unscramble scrambled chromatin fibres forming neighbouring chromosome territories. Chromatin loop extrusion can also maintain chromosomes as essentially unknotted and uncatenated with each other even if some knotting and catenation events may occur between sequential rounds of chromatin loop extrusion. The rare knots observed in single cell Hi-C (32) are presumably results of knotting events that occurred after the latest round of chromatin loop extrusion and before cell fixation for Hi-C.

After this work was completed, a very similar idea of chromatin loop extrusion involvement in chromatin unknotting and decatenation was proposed in a preprint by Orlandini et al. (33).

## MATERIAL AND METHODS

### Description of the model

Molecular dynamics simulations presented in Figure 1 and 2 were performed using Extensible Simulation Package for Research on Soft Matter (ESPResSo) (34) and those presented in Figure 3 were performed using a general-purpose particle simulation toolkit HOOMD-blue (35-37). Chromatin chains were modelled as standard, beaded self-avoiding chains that were composed of 133 beads for the chains presented in Fig. 1 and of 400 beads for the construct presented in Figure 2. In these coarse grained simulations each bead is assumed to have 10 nm diameter and corresponds to a portion of chromatin fibre containing 400 bp (31). The large simulated system presented in Figure 3 consisted of 8 chains, each with the length corresponding to 4000 beads placed after each other and thus representing 1.6 Mb large chromatin portions. Non-standard features built into our models included active movements of cohesin handcuffs (Figure 1 and 2) or progressing shifting of the points of close contacts between contacting chains (Figure 3). Cohesin handcuffs embracing modelled chromatin fibres were composed of 2 small rings (each with 7 beads). These rings were connected with each other using two additional bonds involving two neighbouring beads of each ring. The cohesin rings were advancing along embraced chains by a walking mechanism where bonds with the spring-like potential and the starting length of up to 2 (where 1 is the diameter of beads) were dynamically formed between three non-consecutive beads in each cohesin ring with sequentially called beads within embraced chromatin fibres. As the spring-like bonds were very strong (*K* = 20 *k*B*T*) and their rest length was set to 1, they were rapidly shrinking, which resulted in the advancement of cohesin rings along embraced chromatin fibres. Once these spring-like bonds have shrunk to about their rest length, they were replaced with extended bonds connecting cohesin rings with the consecutive beds of modelled chromatin fibres. Due to the tightness of cohesin rings and their excluded volume potential as well as due to the excluded volume potential of beads forming chromatin fibres, the advancing cohesin rings were effectively pushing forward all encountered entanglements on modelled chromatin fibres. These entanglements were however free to untangle when they were pushed towards TAD borders, where short sections of chains without excluded volume mimicked the presence of TopIIB at TAD borders (19). In the simulated system composed of 8 large chains, the process of chromatin loop extrusion was modelled in a simpler way. Initially, a small loop was generated by establishing a bond with a strong the spring-like potential (*K* = 100 *k*B*T*) between two non-consecutive beads. Once the two beads connected with that spring-like bond were brought close together, the spring-like bond was moved to bring together the next pair of beads and thus increasing the size of the formed loop with each move.

### Voronoi space tessellation

The extent of mutual intermingling of separate chains was evaluated by using the Voronoi space tessellation approach (26). Using this approach the sphere containing 8 chains with 4000 beads each is dissected creating a polyhedron around each of the polymer beads. These polyhedrons represent regions in which every point is closer to the centre of a given bead than to the centre of any other bead. Each of the polyhedrons is created with four or more facets with a given surface area. The total area of the Voronoi facets separating beads belonging to different chains represent the total interfacial area (Figure 3). The interfacial area increases with increasing extent of mutual intermingling between different chains. The tessellation was constructed in a non-periodic space with a spherical wall. The coordinates of Voronoi vertices, indices, facets with respective areas of the interfaces were obtained by using VORO++ software (26). The visualizations were made using POVRay.

### Knots identification

To determine and present the knot types formed after topological equilibration of 8 large rings, we entered xyz coordinates of each individual ring into Knoto-ID (38) to simplify trajectories of formed knots, while maintaining their original topology.

## Acknowledgements

We thank Dorothy Buck and Sean D. Colloms for their comments and suggestions concerning this work.

## References

1. Bates AD & Maxwell A (2005) DNA Topology (Oxford University Press, Oxford) p 198.

2. Bjorkegren C & Baranello L (2018) DNA Supercoiling, Topoisomerases, and Cohesin: Partners in Regulating Chromatin Architecture? International journal of molecular sciences 19(3).

3. Valdes A, Segura J, Dyson S, Martinez-Garcia B, & Roca J (2018) DNA knots occur in intracellular chromatin. Nucleic acids research 46(2):650–660.

4. O’Donnol D, Stasiak A, & Buck D (2018) Two convergent pathways of DNA knotting in replicating DNA molecules as revealed by theta-curve analysis. Nucleic acids research.

5. Portugal J & Rodriguez-Campos A (1996) T7 RNA polymerase cannot transcribe through a highly knotted DNA template. Nucleic acids research 24:4890–4894.

6. Deibler RW, Mann JK, Sumners DW, & Zechiedrich L (2007) Hin-mediated DNA knotting and recombining promote replicon dysfunction and mutation. BMC Mol Biol 8(1):44.

7. Lieberman-Aiden E, et al. (2009) Comprehensive mapping of long-range interactions reveals folding principles of the human genome. Science (New York, N.Y.) 326(5950):289–293.

8. Grosberg A, Rabin Y, Havlin S, & Neer A (1993) Crumpled globule model of the three-dimensional structure of DNA. Europhysics Letters 23(5):373.

9. Mirny LA (2011) The fractal globule as a model of chromatin architecture in the cell. Chromosome research: an international journal on the molecular, supramolecular and evolutionary aspects of chromosome biology 19(1):37–51.

10. Yan J, Magnasco MO, & Marko JF (1999) A kinetic proofreading mechanism for disentanglement of DNA by topoisomerases. Nature 401(6756):932–935.

11. Buck GR & Zechiedrich EL (2004) DNA disentangling by type-2 topoisomerases. Journal of molecular biology 340(5):933–939.

12. Rao SS, et al. (2014) A 3D map of the human genome at kilobase resolution reveals principles of chromatin looping. Cell 159(7):1665–1680.

13. Sanborn AL, et al. (2015) Chromatin extrusion explains key features of loop and domain formation in wild-type and engineered genomes. Proc Natl Acad Sci U S A 112(47):E6456–6465.

14. Fudenberg G, et al. (2016) Formation of chromosomal domains by loop extrusion. Cell reports 15(9):2038–2049.

15. Schwarzer W, et al. (2017) Two independent modes of chromatin organization revealed by cohesin removal. Nature 551(7678):51–56.

16. Fudenberg G, Abdennur N, Imakaev M, Goloborodko A, & Mirny LA (2017) Emerging evidence of chromosome folding by loop extrusion. Cold Spring Harbor symposia on quantitative biology 82:45–55.

17. Dixon JR, et al. (2012) Topological domains in mammalian genomes identified by analysis of chromatin interactions. Nature 485(7398):376–380.

18. Nora EP, et al. (2012) Spatial partitioning of the regulatory landscape of the X-inactivation centre. Nature 485(7398):381–385.

19. Uuskula-Reimand L, et al. (2016) Topoisomerase II beta interacts with cohesin and CTCF at topological domain borders. Genome biology 17(1):182.

20. Canela A, et al. (2017) Genome Organization Drives Chromosome Fragility. Cell 170(3):507-521.e518.

21. Hansen AS, Pustova I, Cattoglio C, Tjian R, & Darzacq X (2017) CTCF and cohesin regulate chromatin loop stability with distinct dynamics. eLife 6.

22. Spengler SJ, Stasiak A, & Cozzarelli NR (1985) The Stereostructure of Knots and Catenanes Produced by Phage-Lambda Integrative Recombination - Implications for Mechanism and DNA-Structure. Cell 42(1):325–334.

23. Meaburn KJ & Misteli T (2007) Cell biology: chromosome territories. Nature 445(7126):379–781.

24. Sugawara T & Kimura A (2017) Physical properties of the chromosomes and implications for development. Development, growth & differentiation 59(5):405–414.

25. Bolzer A, et al. (2005) Three-dimensional maps of all chromosomes in human male fibroblast nuclei and prometaphase rosettes. PLoS biology 3(5):826–842.

26. Rycroft CH, Grest GS, Landry JW, & Bazant MZ (2006) Analysis of granular flow in a pebble-bed nuclear reactor. Physical review. E, Statistical, nonlinear, and soft matter physics 74(2 Pt 1):021306.

27. Branco MR & Pombo A (2006) Intermingling of chromosome territories in interphase suggests role in translocations and transcription-dependent associations. PLoS biology 4(5):e138.

28. Dorier J & Stasiak A (2009) Topological origins of chromosomal territories. Nucleic acids research 37(19):6316–6322.

29. Racko D, Benedetti F, Dorier J, & Stasiak A (2018) Transcription-induced supercoiling as the driving force of chromatin loop extrusion during formation of TADs in interphase chromosomes. Nucleic acids research 46(4):1648–1660.

30. Alipour E & Marko JF (2012) Self-organization of domain structures by DNA-loop-extruding enzymes. Nucleic acids research 40(22):11202–11212.

31. Goloborodko A, Imakaev MV, Marko JF, & Mirny L (2016) Compaction and segregation of sister chromatids via active loop extrusion. eLife 5.

32. Siebert J, et al. (2017) Are There Knots in Chromosomes? Polymers 9(8):317.

33. Orlandini E, Marenduzzo D, & Michieletto D (2018) Synergy of SMC and topoisomerase creates a universal pathway to simplify genome topology. arXiv:https://arxiv.org/pdf/1809.01267.pdf.

34. Limbach HJ, Arnold A, Mann BA, & Holm C (2006) ESPResSo - an extensible simulation package for research on soft matter systems. Comp. Phys. Comm. 174(9):704–727.

35. Anderson JA, Lorenz CD, & Travesset A (2008) General purpose molecular dynamics simulations fully implemented on graphics processing units. Journal of Computational Physics 227:5342–5359.

36. Glaser J, et al. (2015) Strong scaling of general-purpose molecular dynamics simulations on GPUs. Computer Physics Communications 192:97–107.

37. Reith D, Mirny L, & Virnau P (2011) GPU based molecular dynamics simulations of polymer rings in concentrated solution: structure and scaling. Progress of Theoretical Physics Supplement (191):135–145.

38. Dorier J, Goundaroulis D, Benedetti F, & Stasiak A (2018) Knoto-ID: a tool to study the entanglement of open protein chains using the concept of knotoids. Bioinformatics (Oxford, England):10.1093/bioinformatics/bty1365.

